# Extinction Training Suppresses Activity of Fear Memory Ensembles Across the Hippocampus and Alters Transcriptomes of Fear-Encoding Cells

**DOI:** 10.1101/2023.12.31.573787

**Authors:** Alfredo Zuniga, Jiawei Han, Isaac Miller-Crews, Laura A. Agee, Hans A. Hofmann, Michael R. Drew

**Affiliations:** Department of Neuroscience, The University of Texas at Austin; Institute for Neuroscience, The University of Texas at Austin; Department of Integrative Biology, The University of Texas at Austin; Interdisciplinary Life Sciences Graduate Programs, The University of Texas at Austin

**Author notes:** **Corresponding authors:** Dr. Michael R. Drew, Department of Neuroscience, The University of Texas at Austin, 1 University Station, Stop C7000, Austin, TX 78712-0805, Dr. Hans A. Hofmann, Department of Integrative Biology, The University of Texas at Austin, 2415 Speedway – C0930, Austin, TX 78745, USA. These authors contributed equally to this work (Alfredo Zuniga, Jiawei Han). Dr. Alfredo Zuniga, Department of Neuroscience, The College of Wooster, 1189 Beall Ave, Wooster OH, 44691.

## Abstract

Contextual fear conditioning has been shown to activate a set of “fear ensemble” cells in the hippocampal dentate gyrus (DG) whose reactivation is necessary and sufficient for expression of contextual fear. We previously demonstrated that extinction learning suppresses reactivation of these fear ensemble cells and activates a competing set of DG cells – the “extinction ensemble.” Here, we tested whether extinction was sufficient to suppress reactivation in other regions and used single nucleus RNA sequencing (snRNA-seq) of cells in the dorsal dentate gyrus to examine how extinction affects the transcriptomic activity of fear ensemble and fear recall-activated cells. Our results confirm the suppressive effects of extinction in the dorsal and ventral dentate gyrus and demonstrate that this same effect extends to fear ensemble cells located in the dorsal CA1. Interestingly, the extinction-induced suppression of fear ensemble activity was not detected in ventral CA1. Our snRNA-seq analysis demonstrates that extinction training markedly changes transcription patterns in fear ensemble cells and that cells activated during recall of fear and recall of extinction have distinct transcriptomic profiles. Together, our results indicate that extinction training suppresses a broad portion of the fear ensemble in the hippocampus, and this suppression is accompanied by changes in the transcriptomes of fear ensemble cells and the emergence of a transcriptionally unique extinction ensemble.

## INTRODUCTION

Although fear-based learning is important for survival, learned fear that is excessive and/or overgeneralized – as is seen in clinical phobias or post-traumatic stress disorder – can be debilitating. The front-line behavioral methods for treating such fear-related disorders are based on extinction. In these treatments, patients are repeatedly exposed to the feared stimuli in a safe context [1]. This will ideally produce fear extinction, a lasting decrease in the behavioral response to the feared stimuli and a critical component of adaptive and context-appropriate behavior [2] Unfortunately, fear often relapses after extinction. For instance, in spontaneous recovery the fear response returns with the passage of time after the conclusion of extinction training [3]. The existence of spontaneous recovery and other forms of relapse imply that, rather than abolishing or otherwise altering the fear memory, extinction training works by inducing formation of a competing “extinction memory” that inhibits expression of the fear memory.

Research on the neural mechanisms of extinction has identified the hippocampus as a crucial structure in this form of learning. Disruption of hippocampal activity both interferes with extinction learning [4–6] and can alter generalization of the extinction memory to other contexts [7–9]. Studies using immediate early gene (IEG)-based trapping approaches to tag cells active in a specific timeframe demonstrate that the hippocampus generates ensemble representations of both contextual fear and contextual fear extinction [6,10,11,61,62]. Neurons active during context fear conditioning (CFC)—the fear ensemble—are reactivated during recall of the memory, and this reactivation is both necessary and sufficient for expression of contextual fear. We recently demonstrated that extinction training suppresses reactivation of the fear ensemble within the hippocampal dentate gyrus (DG) and activates a separate set of cells, the putative extinction ensemble [6]. Whereas artificial stimulation of the fear ensemble increases fear, stimulation of this extinction ensemble reduces it.

In the present study, we investigate the effects of extinction training on IEG activity and gene expression in the hippocampal fear ensemble. We first set out to evaluate the generality of the effects of extinction on the fear ensemble. In Lacagnina et al [6], extinction was shown to suppress expression of the IEG Arc in DG fear neurons; other regions and activity markers were not evaluated. Here, we asked whether the suppressive effect of extinction was present beyond the DG, whether this effect was present across the dorsoventral axis of the hippocampus, and whether it could be observed with another canonical IEG, c-Fos. A second objective was to characterize the effects of extinction training on the transcriptomes of fear acquisition neurons. Our results demonstrate that extinction training causes a robust suppression of both Arc and c-Fos expression in the hippocampal fear ensemble throughout the dorsal and ventral DG and the dorsal CA1. This suppression is accompanied by profound transcriptional changes in the fear ensemble, providing novel insights into the molecular mechanisms through which extinction suppresses learned fear.

## MATERIALS & METHODS

### Animals

Adult male and female ArcCreERT2::eYFP mice (12-15 weeks) were used for all experiments. For experiment 1 (Fig. 1-3), 12 mice were used (6 per group). An additional 6 mice were used for experiment 2 (Fig. 4,5). Mice were generated by breeding pairs gifted by Dr. Christine Denny [11]. Briefly, heterozygous (+/-) ArcCreERT2::(+/+) R26R-STOP-floxed-eYFP mice were crossed with homozygous (-/-) ArcCreERT2::(+/+) R26R-STOP-floxed-eYFP mice. These crosses generated mice that were heterozygous (+/-) for the ArcCreERT2 allele and homozygous (+/+) for the eYFP allele. Mice were housed in groups (4-5 per cage) until 7 d prior to the start of the experiment, at which point they were individually housed in plastic cages with woodchip bedding. Mice were kept on a 12 h light/dark cycle (lights on at 07:00) with *ad libitum* access to food and water. All protocols were reviewed and approved by the University of Texas at Austin Institutional Animal Care and Use Committee.

**Fig. 1.**
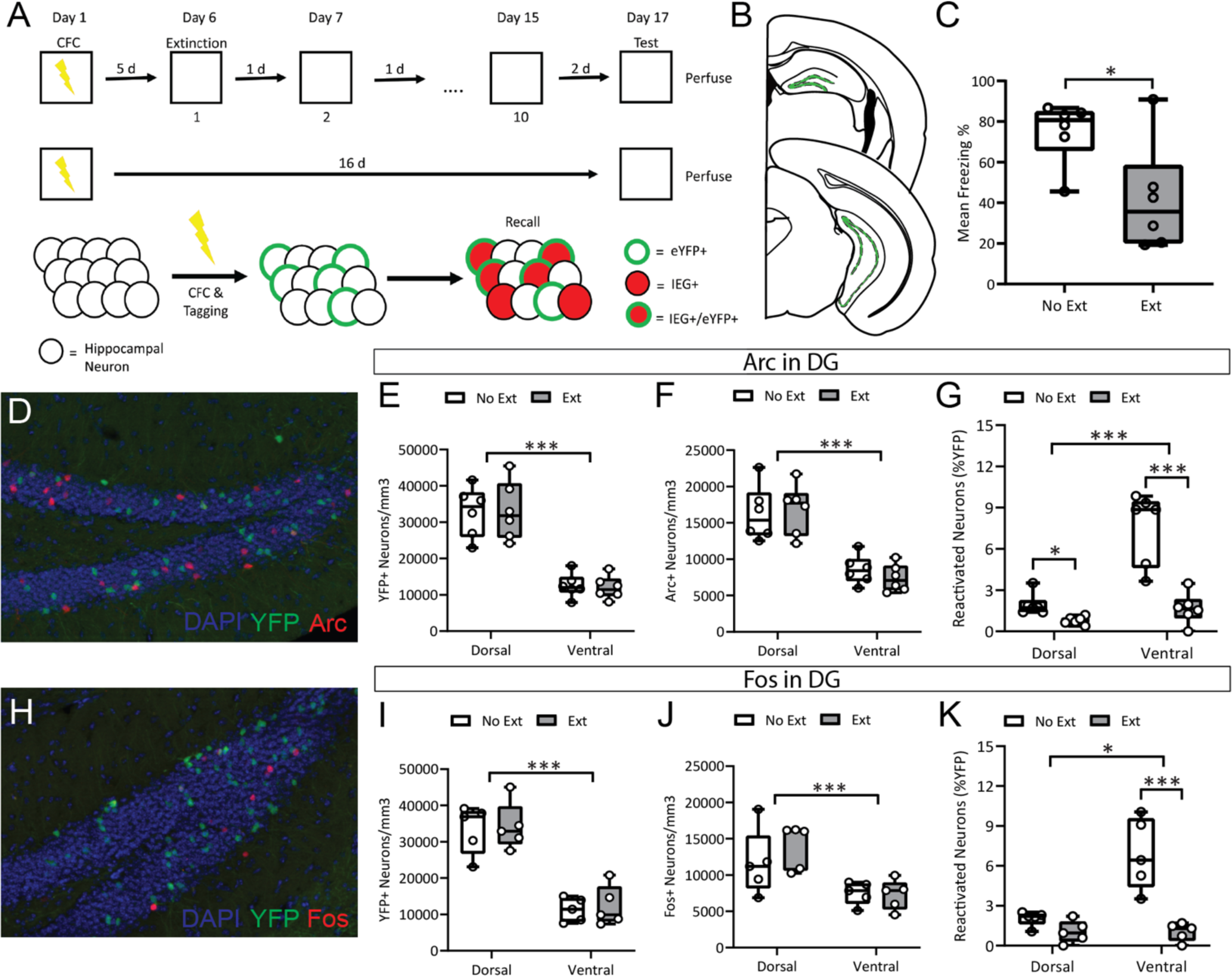
Immediate-early gene activity in the dentate gyrus following extinction or fear recall. **A.** Experimental design. ArcCreERT2::eYFP mice were injected with 4-hydroxytamoxifen immediately following contextual fear conditioning (CFC) to tag CFC activated neurons. Mice then either underwent fear extinction (*Ext group)* or were left undisturbed (*No Ext group)* before being re-exposed to the CFC context and having their tissue processed for Arc or c-Fos expression. **B.** Coronal slices displaying (in green) representative locations from which dorsal (back) and ventral (front) dentate gyrus (DG) cells counts were taken. **C.** Mice that received extinction training displayed significantly less context freezing on the final testing day.**D.** Representative micrographs of YFP+ and Arc+ immunofluorescence in the DG. Tissue processed for Arc expression (E-G) showed no group differences in YFP+ or Arc+ cells in either the dorsal or ventral DG. **H.** Representative micrographs of YFP+ and Fos+ immunofluorescence in the DG. Tissue processed for c-Fos expression (I-K) similarly showed no difference in overall YFP+ or Fos+ cell counts. In contrast, *No Extinction* mice displayed significantly higher levels of Arc+/YFP+ reactivated cells (G) and Fos+/YFP+ reactivated cells (K) in both the dorsal and ventral DG in comparison to mice in the *Extinction* group. *\* p < 0.05, ** p <*

### Neuronal tagging

Recombination was induced with 4-hydroxytamoxifen (4-OHT, Sigma-Aldrich), as previously reported [6,11]. 4-OHT (10 mg/mL) was dissolved via sonication in a 10% ethanol/90% sunflower seed oil solution. To minimize transportation- and novelty-induced IEG activity, mice were transported to a separate housing room near the conditioning room 24 h prior to fear conditioning and 4-OHT injections. Neuronal tagging was induced by a single intraperitoneal injection of 4-OHT (55 mg/kg) given immediately following the contextual fear conditioning session. Following the 4-OHT injection, mice were placed back in the housing room for 72 h during which they were housed in the dark. Previous studies have shown that housing mice in the dark following 4-OHT administration minimizes non-specific neuronal tagging [11]. At the conclusion of the 72 h, mice were returned to the vivarium with a 12 h light/dark cycle for the duration of the experiment.

### Contextual fear conditioning

All mice were handled once a day for 1-2 min for 3 d prior to the commencement of fear conditioning. One hour prior to the start of the contextual fear conditioning session, mice were transported from the separate housing room (see above) to a holding room adjacent to the behavioral room. Mice were moved individually from their home cage to and from the conditioning room in an opaque container with a clear lid. Transport containers were cleaned between mice with 70% ethanol.

Fear conditioning, extinction, and testing occurred in conditioning chambers (30.5 x 24 x 21cm, Med Associates) constructed from two aluminum side walls, a clear Plexiglass door and ceiling, and a white vinyl back plate. Chambers were contained within larger, sound-attenuating chambers that were continuously lit with an overhead white light throughout all behavioral procedures. All mice were exposed to the same conditioning context, which consisted of a stainless-steel rod floor (36 rods, 8mm apart) scented with 1% acetic acid in the waste tray located below the floor. The fear conditioning protocol consisted of three 2-sec, 0.75 mA scrambled foot shocks delivered via the floor 180, 240 and 300 sec after mice were placed in the chambers. Mice were removed from the conditioning chambers and returned to their home cages 30 sec after the final foot shock. All behavior was recorded at 30 frames per second using a near-infrared camera mounted to the door of the chamber, and freezing behavior was scored using VideoFreeze Software (Med Associates).

### Extinction and testing

Extinction sessions consisted of a 5-min exposure to the conditioning context without the presentation of a shock. Mice in the *Extinction* group were exposed to extinction training once a day for 10 days. Mice in the *No Extinction* group were left undisturbed in their home cages (in the vivarium) during these sessions. 72 h after the last extinction session, all mice were exposed to a final 5 min CFC test in the conditioning context.

### Tissue collection and processing

Mice were euthanized following the conclusion of the final behavioral test, and tissue was collected in one of two ways depending on whether tissue was to be processed for immunohistochemistry or single nuclei RNA sequencing (snRNA-seq). For immunohistochemistry, mice were anesthetized with ketamine/xylazine (150/15 mg/kg) 90 minutes after the end of behavioral testing and transcardially perfused with 1X PBS followed by 4% paraformaldehyde in 1X PBS. Brains were extracted and fixed in 4% PFA, then immersed in 20-30% sucrose for 48 h. Finally, brains were cryo-sectioned at 35 µm (Leica) and stored in cryo-protectant at -20 degrees C until staining. For snRNA-seq, mice were anesthetized with 5% isoflurane and then rapidly decapitated 30 min after the final behavioral test. Brains were extracted and flash frozen in OCT cryostat embedding medium (Sakura Finetek) and then stored at -80 degrees C until further processing.

### Immunohistochemistry and imaging

Eight to ten sections (per animal) encompassing the dorsal and ventral hippocampus (Bregma -1.7 to -3.64mm) were washed in 1X PBS (3 x 10 min) and blocked in 5% normal donkey serum (NDS) in 1X PBS with 0.25% Triton X-100 (PBST) for 90 min. Sections labeled for only one IEG were incubated with 1:2000 chicken anti-GFP (Aves Labs) and either 1:2000 rabbit anti-Fos (Synaptic Systems) or 1:2000 rabbit anti-Arc (Synaptic Systems) primary antibodies, diluted in 5% NDS overnight at room temperature. Sections were then washed in 1X PBS (3 x10 min), blocked in 5% NDS in PBST for 1 h, and then incubated in 1:1000 biotinylated donkey anti-chicken (Jackson), and 1:1000 donkey anti-rabbit Cy-3 (Jackson) in 5% NDS in PBST for 2 h. Following a final round of PBS washes (3 x 10 min), sections were incubated in 1:500 avidin Cy-2 (Jackson) and 1:1000 DAPI for 1 h at room temperature. Sections labeled for both Arc and c-Fos were incubated in 1:2000 chicken anti-GFP, 1:2000 rabbit anti-Arc and 1:2000 rat anti-Fos. These triple-labeled sections were then incubated with secondary antibodies – 1:1000 biotinylated donkey anti-chicken (Jackson), 1:1000 donkey anti-rabbit Cy-3 (Jackson), and 1:1000 donkey anti-rat Cy-5 (Jackson), followed by incubation with 1:500 avidin Cy-2 (Jackson) and 1:1000 DAPI. All sections were mounted onto slides and cover-slipped with Fluoromount-G (Invitrogen) mounting medium. For each animal, 3-4 sections per region were imaged, counted and analyzed. Sections were imaged at 20X magnification across the z-plane (5-7 images per z-stack) on a Zeiss Axio Imager M2 (Zeiss) and counted using Stereo Investigator software (Mbf).

### Dentate gyrus micro-dissection and nuclei isolation for snRNA-seq

Frozen brains were transferred to a cryostat (Leica) and were allowed to sit in the chamber set to -20 degrees C for 1 hr. Brains were sectioned at 300 µm directly onto glass slides and kept cold on dry ice. The dorsal DG was micro-dissected using a disposable tissue punch tool (Electron Microscopy Sciences; diameter: 0.75 mm) directly into a 1.5ml microcentrifuge tube. Tissue punches were stored at -80 degrees C until further processing. We adapted the protocol by Martelotto [12] for nuclei isolation directly from flash-frozen tissue for use with 10x Genomics Chromium Single Cell 3’ Reagent Kit (cat. pn-1000075; 10x Genomics, USA). The tissue was kept frozen or on ice throughout the protocol to prevent any additional transcriptional activity. First, the tissue was mechanically homogenized to release the nuclei by adding 1.5 ml of chilled Nuclei EZ Lysis Buffer (cat. 3408; Sigma-Aldrich, USA) to the tissue in a 2 mL tube and gently pipetting ∼10-20 times before a 10-min incubation on ice with additional gentle pipetting halfway through and at the end. The nuclei were then filtered through a 70 μm Cell Strainer (cat. 431751; Corning, USA) into a 1.5 mL tube and subsequently centrifuged at 500g for 5 min at 4°C before the supernatant was removed and the nuclei pellet was re-suspended in 500 μl of nuclei wash and resuspension buffer with DAPI supplemented [1x PBS, 2.0% BSA, 0.2 U/µl RNase inhibitor (cat. 03335402001; Roche, Germany), and 10 μg/ml DAPI filtered with a 0.22 μm PES syringe filter (cat. 380111; Nest Scientific, USA)]. Re-suspended nuclei were then filtered through a 40 μm Cell Strainer (cat. 352340; Corning, USA) and a 35 μm Cell Strainer (cat. 352235; Corning, USA). We then used a MA900 cell sorter (Sony Biotechnology Inc., USA) to isolate DAPI-stained single nuclei into 1.5 ml tubes containing PBS-only sheath fluid with the following settings: chill sort; purity: mode; nozzle size: 100 μm; sample pressure: 10 psi. The samples were then concentrated via centrifuge at 500g for five minutes at 4°C, the supernatant was removed, and the samples were re-suspended in 70 μl of nuclei wash and resuspension buffer with DAPI supplemented. The concentration of nuclei was quantified by using two 10 μl aliquots on the Countess 3 Automated Cell Counter (Invitrogen, USA) to aim for a concentration between 700-1200 nuclei/μl.

### Library construction and sequencing

Library construction and sequencing were performed at the Genomic Sequencing and Analysis Facility at The University of Texas at Austin. Briefly, samples were loaded on the Chromium Next GEM Single Cell 3’ reagent kits v3.1 as per manufacturer’s instructions, with ∼10,000 nuclei per pool as a target. Following the guidance of Davis et al. (2019) [13] and Schmid et al. (2021) [14], we estimated that we needed to sequence 7,385 cells/pool from ∼3 samples to detect two cell populations with as few as ∼20 cells/pool with 99% power. The libraries were sequenced on an Illumina HiSeq 2500 platform (Illumina, USA), generating 930 million reads, with 460 million reads per sample on average. All reads had Phred scores above 35 and passed all quality control measures (MultiQC) [15]. All sequence data have been submitted to NCBI (accession number SUB14102351).

### Bioinformatics and Statistics

The data from all behavioral and immunohistochemical experiments were analyzed using the base statistics functions in R alongside statistical functions from the dplyr [19], rstatix [20], and emmeans [21] packages. Specifically, we used t-tests and analysis of variance (ANOVA). Tukey HSD tests were used for post-hoc comparisons. If assumptions of normality were not met, we log-transformed the data. Effect sizes were calculated using the untransformed data and reported as the generalized eta-squared (η_G_^2^) [22] for the full ANOVAs or Cohen’s d for post-hoc pairwise comparisons. For the two pooled snRNA-seq samples, FASTQ files were generated after demultiplexing and subjected to quality control by FAST-QC (Babraham Bioinformatics). Cellranger counts (Cell Ranger 6.1, 10x Genomics) were used to perform alignment, filtering, barcode counting, and UMI counting on the FASTQ files. All reads were mapped to mouse genome, mm10-2020-A. Output files were used to generate Seurat object using Seurat V4.0 [16] in R. Datasets from both samples were pre-processed with the Seurat pipeline with baseline filtering parameters of min.cells = 3 and min.features = 200. The count matrix was filtered to remove empty droplets, doublets and triplets, yielding a total of 8467 nuclei for the No-Extinction sample and 103041 nuclei for the Extinction sample. The data were log-normalized using the NormalizeData function, and highly variable genes were identified using the FindVariableFeatures function. Integrated analysis across the two samples was performed using the Seurat Integration analysis pipeline, generating 24 clusters (FindNeighbors and FindClusters, using first 30 principal components), which were later characterized as 9 putative cell types. Differentially expressed genes (DEGs: p < 0.05, log_2_ fold-change > 0.2) were identified using the R package *limma* Trend [17], and volcano plots were generated using the EnhancedVolcano R package (https://github.com/kevinblighe/EnhancedVolcano). We used the Single Cell Proportion Test (https://github.com/rpolicastro/scProportionTest) with 10,000 permutations to estimate the proportional difference between treatment in cell number across clusters. Functional enrichment analysis was carried out using Metascape [18] to discover enriched Gene Ontology (GO) terms or Kyoto Encyclopedia of Genes and Genomes (KEGG) pathways. Enriched terms were required to include ≥3 candidates, with a p-value ≤0.01 and an enrichment factor ≥1.5. All metadata and protocols/scripts are available on the Texas Data Repository.

## RESULTS

### Behavior

We compared freezing levels at final recall between mice that had undergone extinction (Ext group) and those that had not (No Ext group). As expected, a between subjects t-test showed that mice that underwent extinction training following CFC exhibited significantly less freezing at the final recall test than mice that did not receive extinction training [Fig. 1B; *t*(10) = 2.66, *p* = 0.029].

### Immediate-Early Gene Ensemble Expression

Using this procedure we previously demonstrated that extinction training suppresses reactivation of DG fear acquisition neurons [6]. We sought to replicate this finding and assess its generality across the dorsoventral axis of the various hippocampal subregions. We first used YFP to identify DG neurons that were active during fear conditioning, Arc protein expression to identify neurons active during the test, and overlap of both markers to identify fear ensemble cells reactivated at final testing (%YFP+ cells expressing Arc). Expression of these markers was assessed in the dorsal and ventral DG, and expression across groups was compared using two-way mixed ANOVAs. We found no significant differences in expression of YFP or Arc between the Ext and No Ext groups in the dorsal or ventral DG (Fig. 1E-F; all p > 0.1). There were, however, significantly more Arc+ [F(1, 10) = 69.08, *p* < 0.0001, η_G_^2^ = 0.724] and YFP+ [F(1, 10) = 146.83, *p* < 0.0001, η_G_^2^ = 0.794] cells in the dorsal than the ventral DG. Importantly, we found that reactivation was reduced in the Ext group compared to No Ext mice [F(1,10) = 23.43, p < 0.001, η_G_^2^ = 0.557] and significantly different between region (dorsal vs. ventral DG) [F(1,10) = 22.877, p < 0.001, η_G_^2^ = 0.504] with an interaction between treatment and region [F(1,10) = 8.499, p = 0.015, η_G_^2^ = 0.376] (Fig. 1G). Tukey HSD test comparisons using estimated marginal means determined that the effect of treatment was present in both the dorsal [t = 2.275, p = 0.0354, d = 2.29] and ventral [t = 5.636, p < 0.0001, d = 2.5] DG, with overall fear ensemble reactivation levels being higher in the ventral DG.

Next, we asked whether this suppressive effect of extinction could be observed when a different IEG is used to assess neuronal activation. In a separate set of sections from the same mice, we performed immunohistochemistry against YFP and the IEG c-Fos and quantified expression in the DG. Again, there were no significant differences in overall YFP or c-Fos expression between the No Ext and Ext groups in either the dorsal or ventral DG (Fig. 1H-I), although both YFP [F(1, 8) = 302.19, *p* < 0.0001, η_G_^2^ = 0.824] and c-Fos [F(1, 8) = 31.48, *p* = 0.0005, η_G_^2^ = 0.498] were again more abundant in the dorsal DG. Reactivation (%YFP+ cells expressing c-Fos) was also once again higher in the No Ext group than the Ext group [F(1,8) = 32.47, p < 0.001, η_G_^2^ = 0.625] and higher overall in the ventral DG as compared to the dorsal DG [F(1,8) = 7.801, p = 0.023, η_G_^2^ = 0.453] (Fig. 1J). A Treatment x Region interaction [F(1, 8) = 7.255, *p* = 0.0273, η_G_^2^ = 0.454] was also detected. Pairwise comparisons of estimated marginal means confirmed that reactivation was suppressed by extinction training in the ventral DG [t = 5.886, p < 0.0001, d = 2.96], while the dorsal DG trended towards, but did not quite reach, significance [t = 1.995, p = 0.0634, d = 1.41].

We then asked whether Arc and c-Fos were expressed in the same or different populations of reactivated cells. In a separate set of sections from the same mice as above, we conducted immunohistochemistry against Arc, c-Fos, and YFP and calculated expression in the dorsal DG only. Consistent with the findings above, a one-way ANOVA showed no differences between the Extinction and No Extinction groups in density of YFP+ cells [F(1, 10) = 0.126, p > 0.1, d = 0.205]. A two-way ANOVA against expressed IEGs (Arc+, Fos+, or Arc+/Fos+) also found no effect of treatment [F(1, 10) = 0.131, p > 0.1, η_G_^2^ = 0.013], though a significant effect of IEG type was detected with there being more Arc+/c-Fos+ cells than single positive Arc [t = 15.636, p < 0.001, d = 1.48] or c-Fos [t = 7.943, p < 0.001, d = 0.78] cells (Fig 2C). This is likely due to the fact that expression of both Arc and c-Fos are triggered by neural firing and, therefore, are more likely to be expressed together. There were also more cells single positive for Arc than c-Fos [t = 7.694, p < 0.001, d = 0.681]. Consistent with previous findings, there was significantly more reactivation of YFP+ cells in the No Extinction group as compared to the Extinction group [F(1,10) = 9.275, p = 0.012, η_G_^2^ = 0.263]. An significant effect of IEG type on reactivation level [F(2,20) = 46.576, p < 0.001, η_G_^2^ = 0.742] and an interaction between Treatment and IEG overlap type (Arc+/YFP+, Fos+/YFP+, or Arc+/Fos+/YFP+) were also detected [F(2, 20) = 11.49, *p* < 0.001, η_G_^2^ = 0.415]. Post-hoc pairwise comparisons showed that the suppressive effect of extinction was observed in YFP+ cells co-expressing Arc and c-Fos [t = 5.641, p < 0.001, d = 2.38] but not in YFP+ cells expressing only Arc [t = 0.234, p > 0.1, d = 0.153] or c-Fos [t = 0.214, p > 0.1, d = 0.209] (Fig 2D). Thus, extinction training suppresses co-expression of Arc and c-Fos in YFP-tagged fear acquisition cells.

**Fig. 2.**
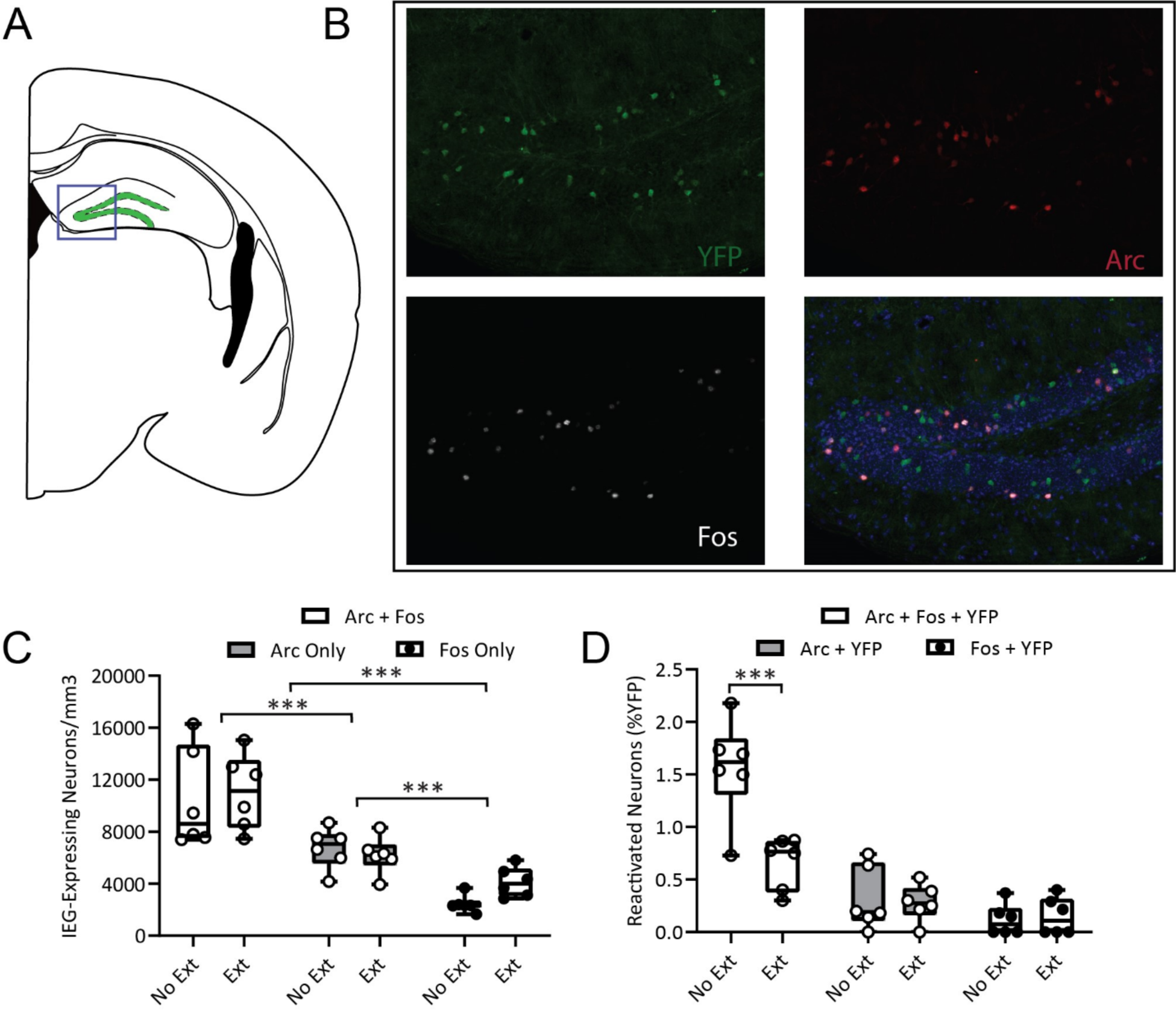
Arc and c-Fos co-expression in the dorsal DG following fear and extinction recall. **A.** Coronal section displaying the representative location of imaging and cell counts. **B.** Representative micrographs of YFP+, Arc+, and Fos+ immunofluorescence in tissue processed for all three proteins. **C.** There was no significant difference between groups in the density of cells expressing Arc+ alone, Fos+ alone, or co-expressing Arc+/Fos+. However, the overall density of cells co-expressing Arc+ and Fos+ was significantly higher than the amount expressing either IEG alone. **D.** Examination of levels of fear tagged (YFP+) cell reactivation found that mice in the *No Extinction* group displayed significantly higher levels of reactivation than mice in the *Extinction* group. This treatment effect was driven by cells that were positive for both Arc and c-Fos alongside YFP. *\*\*\* p < 0.001*

To determine whether the suppressive effect of extinction training on reactivation was preserved further along the hippocampal circuit, we next analyzed expression of YFP and c-Fos in the dorsal and ventral CA1 (Fig 3A,B). A two-way ANOVA revealed a significant difference in YFP counts between the dorsal and ventral CA1 [F(1,7) = 657.33, p < 0.0001, η_G_^2^ = 0.849] and a significant interaction between CA1 region and treatment [F(1,7) = 8.42, p = 0.023, η_G_^2^ = 0.163] which did not survive post-hoc testing [all p > 0.05] (Fig. 3C). No significant differences in c-Fos levels were detected between treatment groups or region and no interaction between variables were detected [all p > 0.1] (Fig. 3D). Comparison of reactivation levels (YFP+/Fos+) across regions and treatments showed a significant effect of treatment [F(1,7) = 29.194, p = 0.001, η_G_^2^ = 0.124], region [F(1,7) = 387.869, p < 0.0001, η_G_^2^ = 0.884], and a significant interaction between the two [F(1,7) = 11.479, p = 0.012, η_G_^2^ = 0.01] (Fig 3E). Post-hoc testing showed that while extinction training significantly decreased reactivation of the fear ensemble in the dorsal CA1 [t = 6.086, p < 0.0001, d = 3.18], no such suppression of reactivation was detected in the ventral CA1 [t = 0.935, p = 0.366, d = 0.68]. Overall, our results suggest that extinction training suppresses reactivation of fear ensembles in the dorsal and ventral DG and the dorsal CA1, but not the ventral CA1.

**Fig. 3.**
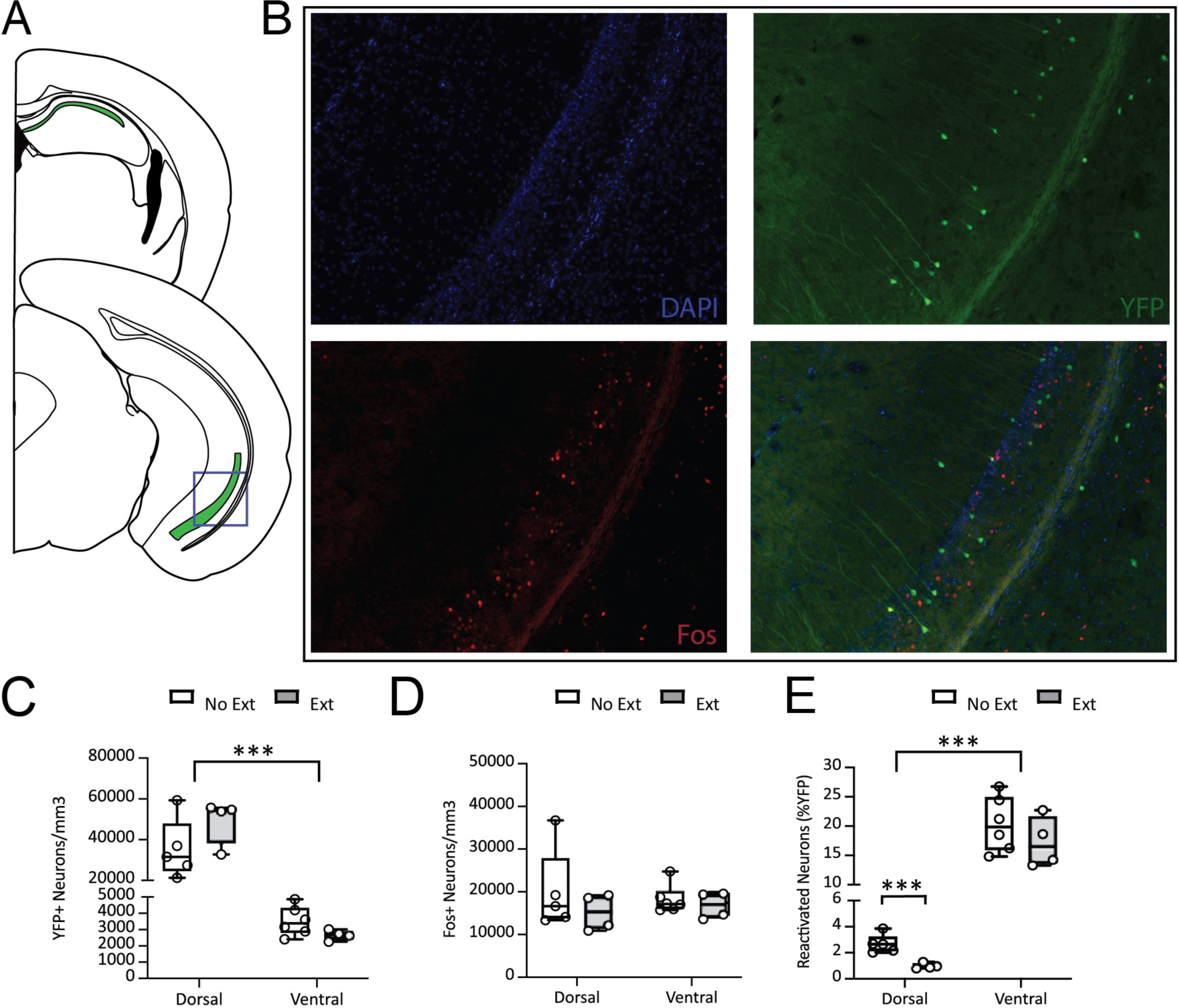
Immediate-early gene activity in the CA1 region of the hippocampus following extinction or fear recall. **A.** Coronal sections displaying (in green) the range and area from which CA1 counts were taken. **B.** Representative images of Fos+ and YFP+ immunofluorescence in the ventral CA1. **C.** There were no differences in CA1 YFP+ cell density between mice in the Ext and No Ext group. In both groups the dorsal CA1 had a higher density of YFP+ cells than the ventral CA1. **D.** There was no effect of extinction training or region of the CA1 on Fos+ cell density. **E.** In the dorsal CA1, mice in the Ext group had significantly fewer fear ensemble cells reactivated (YFP+/Fos+) at final recall as compared to mice in the No Ext group. This effect was not present in the ventral CA1. In both the Ext and No Ext group, more neurons were reactivated at recall in the ventral than dorsal CA1. *\*\*\* p < 0.001*

### snRNA-seq

#### Single-cell transcriptomes reveal canonical cell-types and confirm immunohistochemistry results

We employed snRNA-seq analysis of the dorsal DG as an unbiased and comprehensive approach to survey the transcriptomes of cells belonging to the fear and extinction ensembles. We repeated the behavioral assay as described above (n = 3 per group) and, as before, we observed a significantly less freezing in mice that received extinction training as compared to those that did not [Fig. 4A; t-test: *t*(4) = 10.45, *p* < 0.001]. We then collected and pooled (separately for No Ext and Ext) the dorsal DG nuclei for snRNA-seq. After demultiplexing and filtering, we recovered 19301 nuclei from the two treatment groups (Ext: 10828, No Ext: 8473). To evaluate the cell type-specific response to Fear Extinction/Recall training, we integrated the snRNA-seq datasets from both treatments and then used the Seurat integrative analysis approach revealed 17 clusters that differed in transcriptomic profiles in the integrated dataset (Fig 4C). Based on previously identified hippocampal gene markers [23], these 17 clusters represented nine cell types. As expected, the vast majority of cells were granule cells (Ext: 57.23%; No Ext: 65.22%), although the other canonical cell types (e.g., interneurons, oligodendrocytes, astrocytes, microglia, etc.) were identified as well. In concordance with the immunohistochemistry results above (Fig 1E-J), the number of granule cells that express *c-Fos*, *Arc,* and/or *YFP* mRNA did not appear to differ between treatment groups (Fig 4B1,2), nor did the number of putative extinction ensemble cells (i.e., neurons that expressed *arc* but not *YFP* mRNA) (Fig 4B3). Note that less than 2% of granule cells expressed *c-Fos*, while more than 7% of granule cells expressed *Arc* (a ca. 3.5-fold difference). This is in marked contrast to our immunohistochemistry results, where the difference was only 1.5-fold (see Fig 1F,I). This discrepancy is likely due to the differential kinetics of the *c-Fos* response to acute stimuli: mRNA expression of the gene peaks at approximately 30 min, whereas protein levels peak between 90–120 min [24]. In contrast, hippocampal *Arc* mRNA and protein levels after a simulus exhibit slower and much more correlated kinetics [25].

**Fig. 4.**
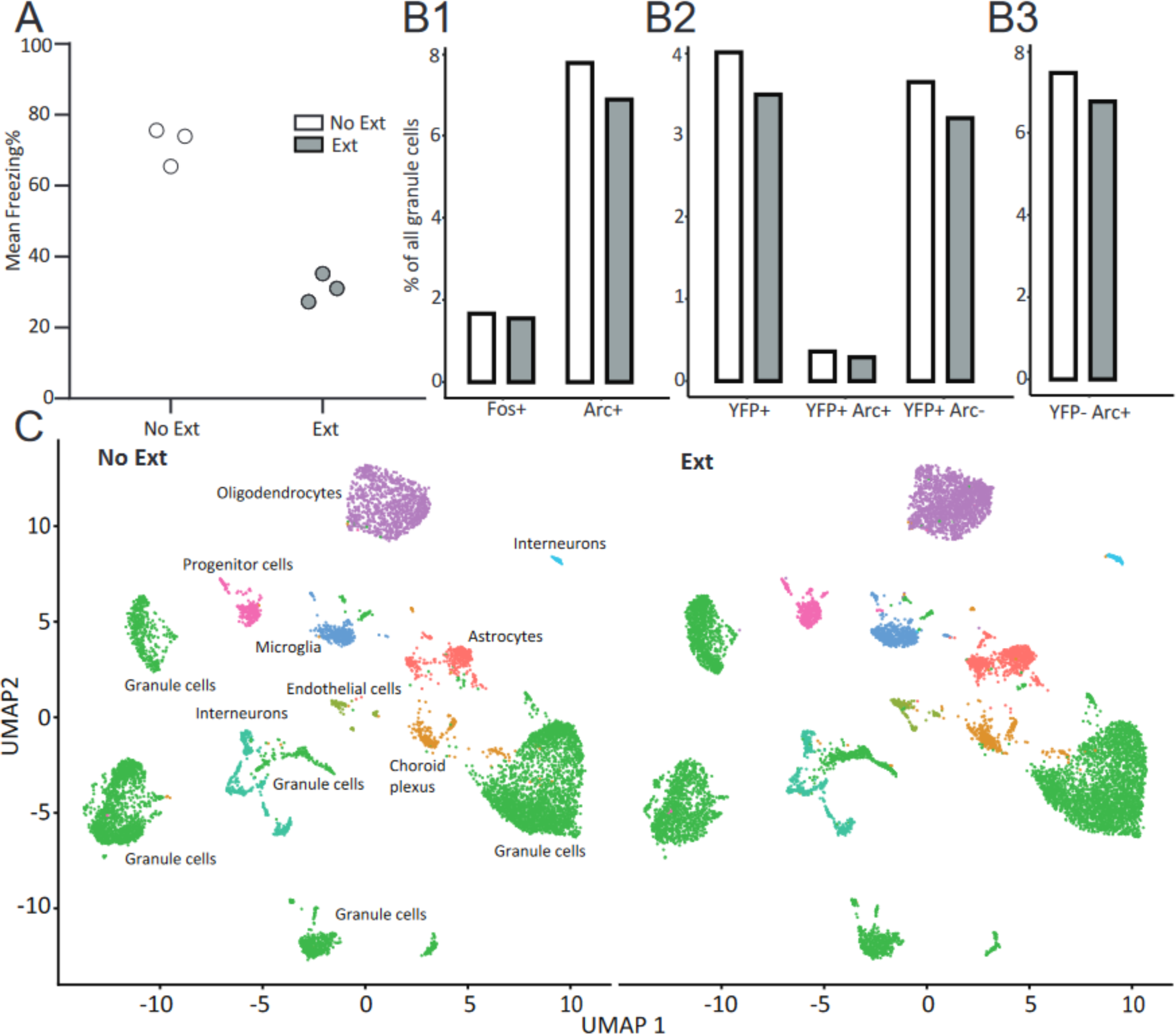
snRNA-seq characterization of dorsal DG in mice after fear extinction training and in control mice. **A.** Behavior test on two groups of animals (Ext: n = 3, No Ext: n =3). **B.** The fraction (%) of granule cells expressing the mRNA for immediate-early genes *c-Fos* or *Arc* (B1); *YFP* only (i.e., fear ensemble cells) or *YFP* and *Arc* (reactivated fear ensemble cells) (B2); and *Arc* in the absence of *YFP* (i.e., putative extinction ensemble cells; B3). **C.** UMAP plots for snRNA-seq data from No Ext and Ext treatments with annotated cell types.

#### Cells activated by extinction training exhibit specific transcriptomic signatures

We have previously shown that extinction training not only suppresses the reactivation of fear ensemble cells but also activates a second set of cells, the putative extinction ensemble [6]. However, little is known about the molecular identity of extinction ensemble cells or how these cells suppress fear ensemble activity. Our dataset enabled us to investigate the transcriptomic response to extinction training at single-cell resolution. We first re-clustered all granule cells and found that both No Ext and Ext groups contained a similar inventory of these cells (412 vs. 419 cells, respectively; Fig 5A). Next, we compared the overall gene expression profiles of the two treatments for those granule cells that expressed *Arc* in the absence of *YFP* mRNA, a cell population that might contain extinction ensemble neurons. We found 1,729 DEGs in these putative “extinction neurons” between Ext and No-Ext samples (Fig 5B; Supplementary Table 1), which were significantly enriched for genes associated with neurodegenerative diseases, among other functions (Supplementary Figure 1A). Interestingly, *Apoe*, a gene that encodes apolipoprotein E, was strongly overexpressed in the Ext group, consistent with previous research showing that *Apoe* dysfunction impairs fear extinction [26,27].

**Fig. 5.**
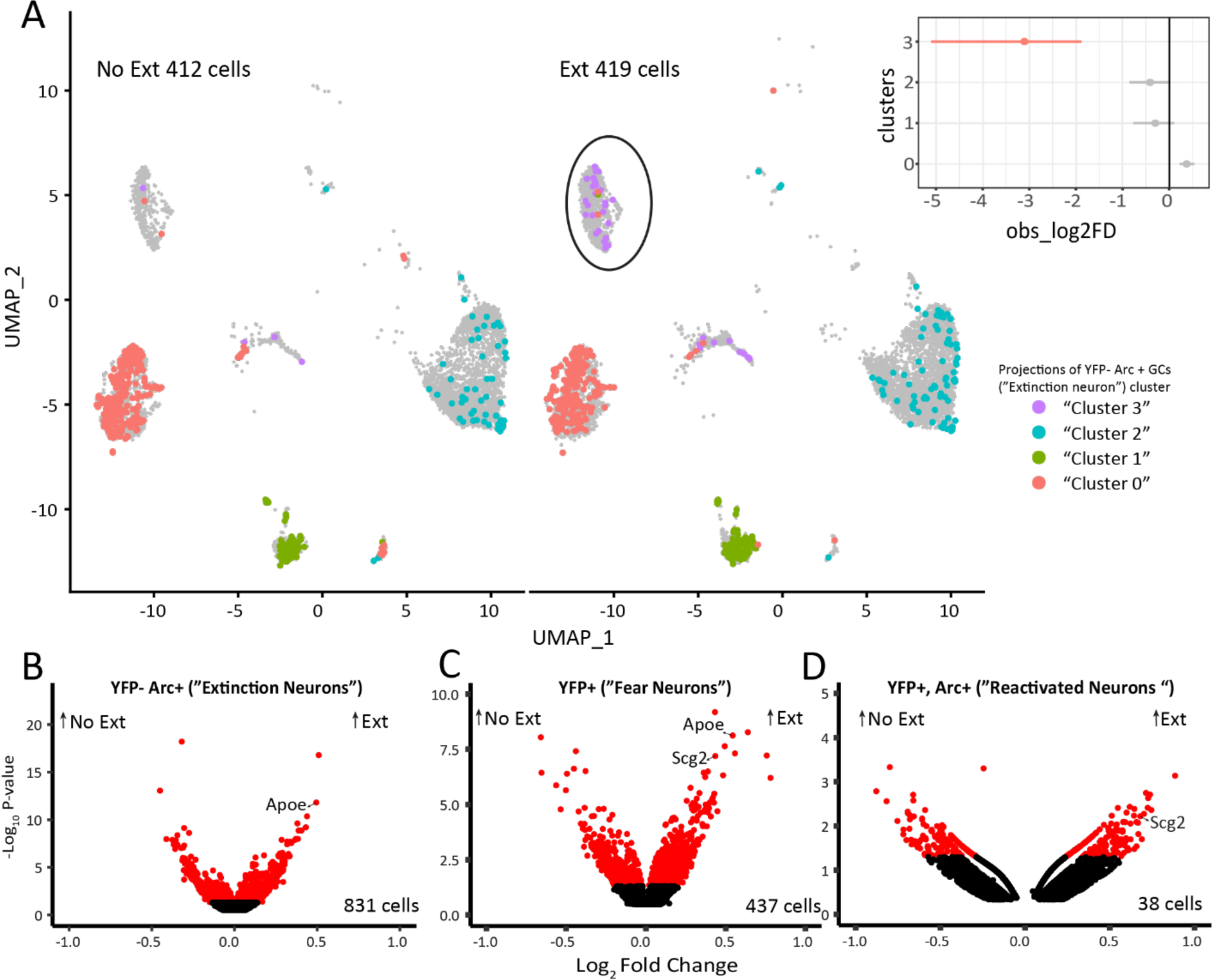
Single nuclei transcription analysis of dorsal dentate gyrus granule cells. **A.** UMAP plots show clustering of granule cells for No Ext and Ext treatments. The putative extinction ensemble cells (i.e., cells that express *Arc* in the absence of *YFP*) are indicated by color according to their cluster identity. Note that “cluster 3” cells (indicated by a circle) are almost exclusively derived from the Ext treatment. Inset shows the relative differences in cell proportions for each cluster between the Extinction and No Extinction groups based on n = 10,000 permutations with Cluster 3 (red) being the only cell cluster with an FDR-adjusted p-value < 0.05 and mean log_2_-fold enrichment > 1. **B.** Volcano plot comparing gene expression of putative extinction ensemble cells in both treatments. DEGs are shown in red (adjusted p-value ≤0.05). **C.** Volcano plot of *YFP* expressing neurons (i.e., fear ensemble cells). **D.** Volcano plot of neurons that co-express *YFP* and *Arc* (i.e., reactivated fear ensemble cells). Specific candidate genes are indicated.

We then asked whether we could identify a unique neuron subtype in the Extinction treatment that could be indicative of a fear extinction ensemble. To achieve this, we extracted putative extinction ensemble cells from both groups (Ext: 419; No Ext: 412) and re-clustered them. Strikingly, this analysis revealed a distinct cluster (cluster 3) of the putative extinction ensemble (Fig 5A inset): 35 of 39 (90%) of the cells in this cluster came from mice in the Ext group (FDR-adjusted p-value < 0.05 and mean log_2_-fold enrichment > 1). None of the other clusters of extinction-activated cells differed significantly between the experimental groups. Because of the 9:1 ratio of cells between treatments in this cluster, it was unfortunately not possible to reliably identify DEGs for these putative extinction ensemble cells. However, GO analysis revealed that the cluster 3 cells were enriched for genes involved in synaptic functions (Supplementary Figure 1B).

#### Secretogranin II (scg2) is a candidate for mediating the suppression of the contextual fear ensemble cells during extinction training

To gain a deeper understanding of the mechanisms underlying the suppression of fear ensemble cells by extinction training, it is critical to examine the transcriptional heterogeneity associated with extinction training. Due to the mRNA expression kinetics of *c-Fos* [24], only a small number of granule cells expressed this IEG in our snRNA-seq dataset, and even fewer co-expressed *YFP* and *c-Fos*. However, we found that 38 of these granule cells co-expressed *YFP* and *Arc* (Ext: 18; No Ext: 20) in our dataset and identified 369 DEGs in these putative “reactivated fear neurons”, including known targets of *c-Fos* (Fig. 5D; Supplementary Table 2). One such target is *Scg2* [28], which encodes the protein secretogranin II and its peptide metabolites. *Scg2* was highly overexpressed in these putative re-activated fear ensemble cells after fear extinction learning. Although the number of these re-activated fear neurons was small, 12 of 18 (66.7%) of these cells expressed *Scg2* in the Ext animals, while only 25% (5 of 20) of these cells did in the No Ext treatment (χ2 = 6.653, p = 0.0099).

Because this result was close to the statistical detection threshold of our dataset (see Methods), we repeated this analysis in the much larger population of *YFP*-expressing fear ensemble cells from both treatments (Ext: 216; No Ext: 221) and identified 1418 DEGs (Fig. 5C; Supplementary Table 3), among them *Scg2* and *Apoe*. These DEGs were enriched for pathways involved in synaptic organization and plasticity as well as genes involved in neural activity (Supplementary Fig. 1C), consistent with the proposed role of *Scg2* in reorganizing inhibitory synaptic inputs in the hippocampus [28]. Finally, to better understand the treatment-specific impact of *Scg2* on the fear ensemble cells, we compared the gene expression profiles of all *Scg2* expressing fear ensemble cells between the two treatment groups (Ext: n = 108; No Ext: n = 55) and identified 803 DEGs (Supplementary Table 4). In line with the strong co-expression of *Scg2* and *Apoe* in fear acquisition neurons observed above, genes that were significantly over-expressed in the Ext treatment were enriched for Alzheimer’s disease genes among other functions (Supplementary Figure 1D).

## DISCUSSION

In the present study, we first examined activity-dependent cell tagging in ArcCreERT2::eYFP transgenic mice to indelibly label cells activated following CFC to test how extinction training affects the reactivation of fear ensemble cells. Our past research has shown that extinction suppresses fear ensemble reactivation in the DG as a whole [6], and here we first expanded on this finding by examining whether the suppressive effect of extinction extends across the dorsoventral axis of the DG. It has long been theorized that the dorsal and ventral hippocampus serve functionally distinct roles for learning and memory [29,30]. Dorsal hippocampal damage is consistently more disruptive to spatial tasks [31–34]. In contrast, ventral hippocampal damage/inhibition primarily disrupts emotional processes [35–39]. The distinct input/output connectivity [40] and genetic expression profiles [41] of the dorsal versus ventral hippocampus further support the idea of a dorsoventral functional dichotomy. One might expect that extinction of a contextual fear association should preserve activity in cells encoding the spatial information about the context while suppressing activity in cells encoding the emotional association. That we see suppression of the fear ensemble across the dorsoventral axis of the DG following extinction suggests that both dorsal and ventral DG process information related to emotional valence. Importantly, this suppressive effect was observed with both Arc and c-Fos IEGs, lending credence to the idea that the observed changes are robust and functionally significant. It is also worth noting that while the effect of extinction on reactivation was present in both the dorsal and ventral DG, the scale of the effect was larger in the ventral DG. This may suggest a more flexible representation of valence exists within the ventral DG.

Next, we examined whether the suppressive effect of extinction on fear ensemble reactivation would persist downstream of the DG, in the CA1. In contrast to the DG, the dorsal and ventral CA1 responded differentially to extinction, with fear ensembles suppressed by extinction in dorsal CA1 but not ventral CA1. The lack of effect in ventral CA1 is surprising in light of the abundant evidence linking ventral CA1 to emotional processing [42–45]. The lack of extinction-mediated suppression in ventral CA1 may suggest that this region contains a valence-free context representation. Alternatively, ventral CA1 might generate a contextual fear representation that is more durable than those established elsewhere in the hippocampus, such that the ventral CA1 representation persists through extinction. Further experimentation will be needed to evaluate these ideas.

In our second experiment, we used a combination of activity dependent cell tagging in ArcCreERT2::eYFP transgenic mice and snRNA-seq to compare the transcriptomes of DG granule cells. We further stratified our analysis based on whether cells had been activated by fear conditioning, extinction/fear recall, or both. We discovered almost as many *YFP^-^/Arc^+^* cells in the No Ext treatment as in the Ext treatment (412 vs. 419). One might therefore ask which, if any, of these cells are part of an emerging extinction ensemble. We speculate that the cells that constituted the small cluster of *YFP^-^/Arc^+^* expressing granule cells that was almost exclusively limited to the Ext treatment (“cluster 3”) may represent the stabilized extinction ensemble already present in the Ext group 30 minutes after the final behavioral test. Although characterizing these cells in detail will require larger sample sizes, we also suggest that the remaining *YFP^-^/Arc^+^* cells from both treatment groups are in the process of being actively recruited into a newly forming extinction ensemble.

Our analysis of mRNA expression in the wider population of DG granule cells active at the fear learning time point (*YFP+*) showed that in mice that had undergone extinction training, genes related to synaptic reorganization were upregulated in the fear ensemble. One such extinction upregulated gene is *Scg2,* which encodes the protein secretogranin II (Scg2). *Scg2* was of particular interested due to recent work by Yap et al. (2021) [28], in which they demonstrate a *Scg2-*dependent mechanism for reorganizing interneuronal inhibitory inputs to hippocampal cells activated by environmental stimuli. In this work, Yap and colleagues (2021) found that CA1 principal cells activated after exposure to a novel environment subsequently displayed higher sensitivity to inhibition from neighboring parvalbumin-expressing interneurons and lower sensitivity to inhibition from cholecystokinin-expressing interneurons. This effect was dependent on *Scg2* expression within the affected CA1 principal neurons, demonstrating a role for *Scg2* in modulating the strength of inhibitory connections onto hippocampal neurons in an experience-dependent manner. The observed increase in *Scg2* expression in the fear ensemble of extinguished mice suggests that *Scg2* may play a similar role in strengthening inhibitory inputs to the fear ensemble following extinction. Notably, we also observed overexpression of the *Scg2* gene in the sparse number of DG fear ensemble cells that were reactivated in the Ext group. This suggests that *Scg2-*mediated inhibitory strengthening remains ongoing throughout the extinction process even in extinction-resistant fear ensemble cells. Finally, targeted comparison of Scg2-expressing granule cells between treatment groups revealed that in the Ext treatment, Alzheimer’s disease genes showed increased expression. This finding may suggest shared mechanisms between Scg2-mediated fear extinction and neurodegenerative diseases. In line with this idea, a number of studies have demonstrated impaired fear extinction in both humans with [46] and rodent models of Alzheimer’s disease [47–49].

Another notable mRNA product that was upregulated in our extinction group was *Apoe.* We observed *Apoe* upregulation in both the fear ensemble (*YFP+* cells) and the extinction ensemble (*YFP-, Arc+* cells) of extinction-trained mice. *Apoe* is translated into apolipoprotein E (Apoe), a lipoprotein which – along with ApoJ – is one of the primary mediators of cholesterol transport in the central nervous system [50–52]. There is ample evidence that Apoe serves an important role in fear extinction learning. For example, mice that received targeted replacement of the endogenous mouse *APOE* gene sequence for the altered human sequence encoding for the apoE2 isoform allele – but not the apoE4 or apoE3 isoforms – fail to show extinction after fear conditioning [26,27]. Additionally, military veterans who are carriers of the apoE2 isoform are significantly more susceptible to PTSD compared to those who produce other Apoe isoforms [27,53,54]. The biological mechanisms which underlie this impairment in extinction learning in individuals with the apoE2 producing genotype remain unknown, but might relate to the unusually low binding affinity of apoE2 to low-density lipoprotein receptors [55,56]. Low-density lipoprotein receptor knockout consistently display learning and memory impairments on a variety of spatial tasks [57–59] and decreased neurogenesis and synaptic bouton density within the hippocampus [60]. The failure of apoE2 to properly bind to and activate these receptors may result in similar deficiencies.

## CONCLUSION

Together, our findings demonstrate that extinction learning suppresses reactivation of neurons encoding fear associations throughout a variety of hippocampal subregions. This suppression is present across the dorsoventral axis of the hippocampus, with the possible exception of the ventral CA1 fear ensemble. The process of extinction learning also appears to preferentially activate genes that facilitate the development and formation of new synapses, suggesting that the cells recruited in the extinction ensemble actively remap their connectivity in response to extinction learning. Further investigation into the mechanisms driving the formation of an effective extinction ensemble could provide clues as to how dysfunction in the fear learning system arises and how it may be treated.

## ACKNOWLEDGEMENTS

We thank Dr. Nihal Salem for guidance and the members of the Drew and Hofmann labs for discussion. Sequencing was performed by the Genomic Sequencing and Analysis Facility at UT Austin, Center for Biomedical Research Support (RRID# SCR_021713). Computational analyses were performed using the Biomedical Research Computing Facility at UT Austin, Center for Biomedical Research Support (RRID#: SCR_021979). This research was supported by a UT Austin Catalyst seed grant to HAH and MRD, U.S. National Institutes of Health grant R01 MH117426 to MRD, U.S. National Science Foundation grant IOS-1326187 to HAH, a U.S. Department of Justice graduate fellowship to IMC, and UT Austin Graduate School Summer Fellowships to JH and IMC.

## COMEPTING INTEREST STATEMENT

The authors declare no competing interests.

## References

1. Abramowitz JS, Deacon BJ, Whiteside SPH. Exposure Therapy for Anxiety, Second Edition: Principles and Practice. Guilford Publications; 2019. 478 p.

2. Myers KM, Davis M. Behavioral and Neural Analysis of Extinction. Neuron. 2002;36(4):567–84.

3. Rescorla RA. Spontaneous Recovery. Learn Mem. 2004;11(5):501–9.

4. Heldt SA, Stanek L, Chhatwal JP, Ressler KJ. Hippocampus-specific deletion of BDNF in adult mice impairs spatial memory and extinction of aversive memories. Mol Psychiatry. 2007;12(7):656–70.

5. Bernier BE, Lacagnina AF, Ayoub A, Shue F, Zemelman BV, Krasne FB, et al. Dentate Gyrus Contributes to Retrieval as well as Encoding: Evidence from Context Fear Conditioning, Recall, and Extinction. J Neurosci. 2017;37(26):6359–71.

6. Lacagnina AF, Brockway ET, Crovetti CR, Shue F, McCarty MJ, Sattler KP, et al. Distinct hippocampal engrams control extinction and relapse of fear memory. Nat Neurosci. 2019;22(5):753–61.

7. Corcoran KA, Maren S. Hippocampal Inactivation Disrupts Contextual Retrieval of Fear Memory after Extinction. J Neurosci. 2001;21(5):1720–6.

8. Corcoran KA, Maren S. Factors Regulating the Effects of Hippocampal Inactivation on Renewal of Conditional Fear After Extinction. Learn Mem. 2004;11(5):598–603.

9. Hobin JA, Ji J, Maren S. Ventral hippocampal muscimol disrupts context-specific fear memory retrieval after extinction in rats. Hippocampus 2006;16:174–182.

10. Tayler KK, Tanaka KZ, Reijmers LG, Wiltgen BJ. Reactivation of Neural Ensembles during the Retrieval of Recent and Remote Memory. Curr Biol. 2013;23(2):99–106.

11. Denny CA, Kheirbek MA, Alba EL, Tanaka KF, Brachman RA, Laughman KB, et al. Hippocampal Memory Traces Are Differentially Modulated by Experience, Time, and Adult Neurogenesis. Neuron. 2014;83(1):189–201.

12. G Martelotto L. ‘Frankenstein’ protocol for nuclei isolation from fresh and frozen tissue for snRNAseq v2. 2019 May. Available from: https://www.protocols.io/view/frankenstein-protocol-for-nuclei-isolation-from-f-3fkgjkw

13. Davis A, Gao R, Navin NE. SCOPIT: sample size calculations for single-cell sequencing experiments. BMC Bioinformatics. 2019;20(1):566.

14. Schmid KT, Höllbacher B, Cruceanu C, Böttcher A, Lickert H, Binder EB, et al. scPower accelerates and optimizes the design of multi-sample single cell transcriptomic studies. Nat Commun. 2021;12(1):6625.

15. Ewels P, Magnusson M, Lundin S, Käller M. MultiQC: summarize analysis results for multiple tools and samples in a single report. Bioinformatics. 2016;32(19):3047–8.

16. Hao Y, Hao S, Andersen-Nissen E, Mauck WM, Zheng S, Butler A, et al. Integrated analysis of multimodal single-cell data. Cell. 2021;184(13):3573–3587.e29.

17. Ritchie ME, Phipson B, Wu D, Hu Y, Law CW, Shi W, et al. limma powers differential expression analyses for RNA-sequencing and microarray studies. Nucleic Acids Res. 2015;43(7):e47.

18. Zhou Y, Zhou B, Pache L, Chang M, Khodabakhshi AH, Tanaseichuk O, et al. Metascape provides a biologist-oriented resource for the analysis of systems-level datasets. Nat Commun. 2019;10(1):1523.

19. Wickham H, François R, Henry L, Müller K, Vaughan D. dplyr: A Grammar of Data Manipulation. 2023. Available from: https://CRAN.R-project.org/package=dplyr

20. Kassambara A. rstatix: Pipe-Friendly Framework for Basic Statistical Tests. 2023. Available from: https://CRAN.R-project.org/package=rstatix

21. Lenth, Russel V. emmeans: Estimated Marginal Means, aka Least-Squares Means. 2023. Available from: https://CRAN.R-project.org/package=emmeans

22. Olejnik S, Algina J. Generalized eta and omega squared statistics: measures of effect size for some common research designs. Psychol Methods. 2003;8(4):434–47.

23. Cembrowski MS, Wang L, Sugino K, Shields BC, Spruston N. Hipposeq: a comprehensive RNA-seq database of gene expression in hippocampal principal neurons. eLife. 5:e14997.

24. Kovács KJ. Invited review c-Fos as a transcription factor: a stressful (re)view from a functional map. Neurochem Int. 1998;33(4):287–97.

25. Ramírez-Amaya V, Vazdarjanova A, Mikhael D, Rosi S, Worley PF, Barnes CA. Spatial Exploration-Induced Arc mRNA and Protein Expression: Evidence for Selective, Network-Specific Reactivation. J Neurosci. 2005;25(7):1761–8.

26. Olsen RHJ, Agam M, Davis MJ, Raber J. ApoE isoform-dependent deficits in extinction of contextual fear conditioning. Genes Brain Behav. 2012;11(7):806–12.

27. Johnson LA, Zuloaga DG, Bidiman E, Marzulla T, Weber S, Wahbeh H, et al. ApoE2 Exaggerates PTSD-Related Behavioral, Cognitive, and Neuroendocrine Alterations. Neuropsychopharmacology. 2015;40(10):2443–53.

28. Yap EL, Pettit NL, Davis CP, Nagy MA, Harmin DA, Golden E, et al. Bidirectional perisomatic inhibitory plasticity of a Fos neuronal network. Nature. 2021;590(7844):115–21.

29. Moser MB, Moser EI. Functional differentiation in the hippocampus. Hippocampus. 1998;8(6):608– 19.

30. Fanselow MS, Dong HW. Are the Dorsal and Ventral Hippocampus Functionally Distinct Structures? Neuron. 2010;1(65):7–19.

31. Kim JJ, Fanselow MS. Modality-Specific Retrograde Amnesia of Fear. Science. 1992;256(5057):675–7.

32. Moser MB, Moser EI, Forrest E, Andersen P, Morris RG. Spatial learning with a minislab in the dorsal hippocampus. Proc Natl Acad Sci. 199;92(21):9697–701.

33. Ferbinteanu J, McDonald R j. Dorsal/ventral hippocampus, fornix, and conditioned place preference. Hippocampus. 2001;11(2):187–200.

34. Pothuizen HHJ, Zhang WN, Jongen-Relo AL, Feldon J, Yee BK. Dissociation of function between the dorsal and the ventral hippocampus in spatial learning abilities of the rat: a within-subject, within-task comparison of reference and working spatial memory. Eur J Neurosci. 2004;19(3):705–12.

35. Kjelstrup KG, Tuvnes FA, Steffenach HA, Murison R, Moser EI, Moser MB. Reduced fear expression after lesions of the ventral hippocampus. Proc Natl Acad Sci. 2002;99(16):10825–30.

36. Maren S, Holt WG. Hippocampus and Pavlovian Fear Conditioning in Rats: Muscimol Infusions Into the Ventral, but Not Dorsal, Hippocampus Impair the Acquisition of Conditional Freezing to an Auditory Conditional Stimulus. Behav Neurosci. 2004;118(1):97–110.

37. Pentkowski NS, Blanchard DC, Lever C, Litvin Y, Blanchard RJ. Effects of lesions to the dorsal and ventral hippocampus on defensive behaviors in rats. Eur J Neurosci. 2006;23(8):2185–96.

38. Esclassan F, Coutureau E, Di Scala G, Marchand AR. Differential contribution of dorsal and ventral hippocampus to trace and delay fear conditioning. Hippocampus. 2009;19(1):33–44.

39. Felix-Ortiz AC, Beyeler A, Seo C, Leppla CA, Wildes CP, Tye KM. BLA to vHPC Inputs Modulate Anxiety-Related Behaviors. Neuron. 2013;79(4):658–64.

40. Swanson LW, Cowan WM. An autoradiographic study of the organization of the efferet connections of the hippocampal formation in the rat. J Comp Neurol. 1977;172(1):49–84.

41. Dong HW, Swanson LW, Chen L, Fanselow MS, Toga AW. Genomic–anatomic evidence for distinct functional domains in hippocampal field CA1. Proc Natl Acad Sci. 2009;106(28):11794–9.

42. Ciocchi S, Passecker J, Malagon-Vina H, Mikus N, Klausberger T. Selective information routing by ventral hippocampal CA1 projection neurons. Science. 2015;348(6234):560–3.

43. Jimenez JC, Berry JE, Lim SC, Ong SK, Kheirbek MA, Hen R. Contextual fear memory retrieval by correlated ensembles of ventral CA1 neurons. Nat Commun. 2020;11(1):3492.

44. Jimenez JC, Su K, Goldberg AR, Luna VM, Biane JS, Ordek G, et al. Anxiety Cells in a Hippocampal-Hypothalamic Circuit. Neuron. 2018;97(3):670–683.e6.

45. Parfitt GM, Nguyen R, Bang JY, Aqrabawi AJ, Tran MM, Seo DK, et al. Bidirectional Control of Anxiety-Related Behaviors in Mice: Role of Inputs Arising from the Ventral Hippocampus to the Lateral Septum and Medial Prefrontal Cortex. Neuropsychopharmacology. 2017;42(8):1715–28.

46. Nasrouei S, Rattel JA, Liedlgruber M, Marksteiner J, Wilhelm FH. Fear acquisition and extinction deficits in amnestic mild cognitive impairment and early Alzheimer’s disease. Neurobiol Aging. 2020;87:26–34.

47. Hernandez CM, Jackson NL, Hernandez AR, McMahon LL. Impairments in Fear Extinction Memory and Basolateral Amygdala Plasticity in the TgF344-AD Rat Model of Alzheimer’s Disease Are Distinct from Nonpathological Aging. eNeuro. 2022;9(3).

48. Bonardi C, de Pulford F, Jennings D, Pardon MC. A detailed analysis of the early context extinction deficits seen in APPswe/PS1dE9 female mice and their relevance to preclinical Alzheimer’s disease. Behav Brain Res. 2011;222(1):89–97.

49. Rattray I, Scullion GA, Soulby A, Kendall DA, Pardon MC. The occurrence of a deficit in contextual fear extinction in adult amyloid-over-expressing TASTPM mice is independent of the strength of conditioning but can be prevented by mild novel cage stress. Behav Brain Res. 2009;200(1):83–90.

50. Roheim PS, Carey M, Forte T, Vega GL. Apolipoproteins in human cerebrospinal fluid. Proc Natl Acad Sci. 1979;76(9):4646–9.

51. Vance JE, Hayashi H. Formation and function of apolipoprotein E-containing lipoproteins in the nervous system. Biochim Biophys Acta BBA - Mol Cell Biol Lipids. 2010;1801(8):806–18.

52. Pitas RE, Boyles JK, Lee SH, Hui D, Weisgraber KH. Lipoproteins and their receptors in the central nervous system. Characterization of the lipoproteins in cerebrospinal fluid and identification of apolipoprotein B,E(LDL) receptors in the brain. J Biol Chem. 1987;262(29):14352–60.

53. Freeman T, Roca V, Guggenheim F, Kimbrell T, Griffin W s. t. Neuropsychiatric Associations of Apolipoprotein E Alleles in Subjects With Combat-Related Posttraumatic Stress Disorder. J Neuropsychiatry Clin Neurosci. 2005;17(4):541–3.

54. Kim TY, Chung HG, Shin HS, Kim SJ, Choi JH, Chung MY, et al. Apolipoprotein E Gene Polymorphism, Alcohol Use, and Their Interactions in Combat-Related Posttraumatic Stress Disorder. Depress Anxiety. 2013;30(12):1194–201.

55. Weisgraber KH, Innerarity TL, Mahley RW. Abnormal lipoprotein receptor-binding activity of the human E apoprotein due to cysteine-arginine interchange at a single site. J Biol Chem. 1982;257(5):2518–21.

56. Wilson C, Mau T, Weisgraber KH, Wardell MR, Mahley RW, Agard DA. Salt bridge relay triggers defective LDL receptor binding by a mutant apolipoprotein. Structure. 1994;2(8):713–8.

57. Mulder M, Jansen PJ, Janssen BJA, van de Berg WDJ, van der Boom H, Havekes LM, et al. Low-density lipoprotein receptor-knockout mice display impaired spatial memory associated with a decreased synaptic density in the hippocampus. Neurobiol Dis. 2004;16(1):212–9.

58. de Oliveira J, Engel DF, de Paula GC, dos Santos DB, Lopes JB, Farina M, et al. High Cholesterol Diet Exacerbates Blood-Brain Barrier Disruption in LDLr–/– Mice: Impact on Cognitive Function. J Alzheimers Dis. 2020;78(1):97–115.

59. de Oliveira J, Hort MA, Moreira ELG, Glaser V, Ribeiro-do-Valle RM, Prediger RD, et al. Positive correlation between elevated plasma cholesterol levels and cognitive impairments in LDL receptor knockout mice: relevance of cortico-cerebral mitochondrial dysfunction and oxidative stress. Neuroscience. 2011;197:99–106.

60. Mulder M, Koopmans G, Wassink G, Mansouri GA, Simard ML, Havekes LM, et al. LDL receptor deficiency results in decreased cell proliferation and presynaptic bouton density in the murine hippocampus. Neurosci Res. 2007;59(3):251–6.

61. Liu X, Ramirez S, Pang PT, Puryear CB, Govindarajan A, Deisseroth K, Tonegawa S. Optogenetic stimulation of a hippocampal engram activates fear memory recall. Nature, 2012;484(7394), 381–5

62. Ramirez S, Liu X, Lin PA, Suh J, Pignatelli M, Redondo RL, Ryan TJ, Tonegawa S. Creating a False Memory in the Hippocampus. Science. 2013; 341(6144), 387–391

